# Intrinsically disordered nuclear pore proteins show ideal-polymer morphologies and dynamics

**DOI:** 10.1101/571687

**Authors:** Luke K. Davis, Ian J. Ford, Anđela Šarić, Bart W. Hoogenboom

## Abstract

In the nuclear pore complex (NPC), intrinsically disordered nuclear pore proteins (FG nups) form a selective barrier for transport into and out of the cell nucleus, in a way that remains poorly understood. The collective FG nup behaviour has long been conceptualized either as a polymer brush, dominated by entropic and excluded-volume (repulsive) interactions, or as a hydrogel, dominated by cohesive (attractive) interactions between FG nups. Here we compare mesoscale computational simulations with a wide range of experimental data to demonstrate that FG nups are at the crossover point between these two regimes. Specifically, we find that repulsive and attractive interactions are balanced, resulting in morphologies and dynamics that are close to those of ideal polymer chains. We demonstrate that this property of FG nups yields sufficient cohesion to seal the transport barrier, and yet maintains fast dynamics at the molecular scale, permitting the rapid polymer rearrangements needed for transport events.

## INTRODUCTION

Nuclear pore complexes (NPCs) penetrate the nuclear envelope in eukaryotic cells, controlling macromolecular transport between the nucleus and cytoplasm. The NPC enables small (≲ 5 nm in size) molecules to cross the nuclear envelope, but hinders the transport of larger macro-molecules [1–3]. Larger macromolecular cargoes, however, can diffuse through the NPC if they are bound to nuclear transport receptors that have an affinity to the transport barrier. Remarkably, the NPC maintains the transport barrier while thousands of cargoes shuttle in and out of the cell nucleus per second [4]. This transport barrier consists of proteins (nucleoporins) that are rich in phenylalanine (F) and glycine (G) repeats (hence called FG nups, for FG nucleoporins) and that are grafted to the inner wall of the NPC transport channel. These hydrophobic domains grant the FG nups cohesive properties, which may be counterbalanced by electrostatic interactions [5, 6]. FG nups are intrinsically disordered proteins [7], causing them to behave as flexible polymers [8].

For a long time, there have been conflicting views on the dominant interactions pervading FG nup assemblies, and these views continue to define the interpretation of experimental data. On one hand, repulsive interactions have been postulated to result in entropic polymer-brush behaviour of FG nups [9–11]. On the other hand, cohesive (*i.e*., attractive) interactions can cause FG nups to form hydrogels *in vitro* [4, 12, 13]. The combined effects of repulsion, cohesion, grafting, and nanopore confinement lead to a rich landscape of possible polymer behaviour [14–17].

Recently, more nuanced views regarding FG nup interactions have emerged, substantiated by computational models of FG nups at different levels of coarse graining. By defining FG nups down to their specific amino acid composition, such models can relate, e.g., FG nup behaviour to their chemical composition and can explore the effects of mutations [18–20], with the important caveat that the results critically depend on a large number of parameters describing the various sizes, charges, and hydrophobicities of the amino acids. Complementarily, at a coarser (“mesoscale”) level, FG nups have been modelled as homogeneous polymers where the electrostatic, hydrophobic, and hydrophilic interactions are incorporated into one essential interaction parameter [21, 22]. Remarkably, these coarser models have reproduced key functional properties of FG nups, with and without nuclear transport receptors, as observed in experiment [21, 22]. This strongly suggests that NPC transport functionality may be generically understood in terms of mesoscale polymer physics. These and other studies have also indicated that FG nup assemblies display aspects of both entropic and cohesive physical behaviour [16, 21, 22]. Furthermore, there is experimental evidence that isolated intrinsically disordered proteins, including FG nups, show properties that are close to those of “ideal” polymers [23, 24], characterized by a radius of gyration scaling of *R*_*G*_ ∝ *N* ^1/2^, *N* being the number of monomers, where repulsive and attractive interactions balance such that the polymer behaves as if no excluded-volume or other larger-range intramolecular interactions are present [25].

In this paper, we show how such ideal-polymer behaviour compares with experimental data on 27 purified FG domains that are either isolated or grafted at physiological densities (FG nup assemblies) [5, 10, 22, 26, 27]. Specifically, we assess how the experimental morphologies compare with computational predictions on polymer behaviour from assemblies containing repulsive, cohesive, and ideal polymers. Importantly we also investigate how these behaviours translate into the static and dynamic properties of polymers in a context that is relevant to NPC transport functionality.

## METHODS

### A. Molecular Dynamics (MD)

In order to investigate polymer morphology and dynamics we have utilized MD where, following previous work [15, 22, 28], FG nups were modelled as polymers consisting of *N* identical beads with diameter *d* and bond length *r*_0_, both set to 0.76 nm. This choice in polymer model yielded a predicted persistence length of the approximate size of one amino acid (0.38 nm) and agrees with experimental data on FG nups and other disordered polypeptide chains [29, 30], and that approaches the excluded-volume of an FG nup (i.e., the sum of the van der Waals volumes of its amino acids, overestimating it by ∼ 20% on average, see Table S1). Bonds were implemented using a harmonic potential 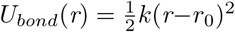 with a spring constant *k* = 500 *k*_*B*_*T/*nm^2^ where *r* is the distance between the centres of two beads.

To capture excluded-volume effects between amino acids and the cohesion arising from attractive –mainly hydrophobic– interactions, we imposed a combined (piecewise) pair potential between polymer beads [31] in which excluded-volume and cohesive interactions were modelled by separate pair potentials so that they can be changed independently. The excluded-volume interaction is the Weeks-Chandler-Andersen (WCA) potential given by

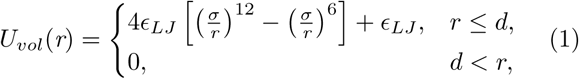

where *ϵ*_*LJ*_ = 500 *k*_*B*_*T* is the interaction strength and 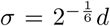 the addition of *ϵ*_*LJ*_ to the potential ensured that *U*_*vol*_(*r* = *d*) = 0.0 *k*_*B*_*T*. The cohesive interaction is based on an infinitely ranged attractive pair potential

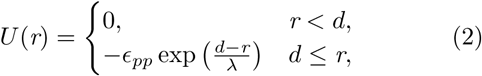

where *ϵ*_*pp*_ is the cohesion strength and *λ* is the decay length [15]. We set *λ* = 0.76 nm and we imposed that no two beads interacted beyond the cutoff distance *r*_*c*_ = 1.52 nm, as it is not possible to have an infinitely-ranged potential in MD. In order to ensure the continuity of the pair potential at *r*_*c*_ we truncated and shifted the potential given by equation 2 using 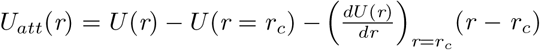 [32] where *U*_*att*_(*r*) is the resulting cohesive pair potential given by

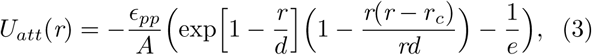

where *e* is Euler’s number and *A* = 1+(*d/l*)^2^ −1*/e*, where *l* = 1.0 nm is the unit of length. This ensured that the minimum of *U*_*att*_(*r* = *d*) = *ϵ*_*pp*_. The total bead-bead pair potential, *U*_*pp*_(*r*), with well depth *ϵ*_*pp*_ is then given as

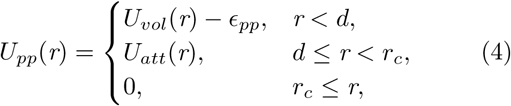

where the minimum of *U*_*vol*_(*r*) is bought down to *ϵ*_*pp*_ to ensure continuity at *r* = *d* (Figure S1a). MD simulations were performed using the LAMMPS package (2016) [33]. We subjected the polymer system to Langevin dynamics at a constant temperature, *T*, by implementing the *NVE* (constant number of beads *N*, constant volume *V*, constant total energy *E*) time integration algorithm in combination with a Langevin thermostat. We performed the simulations with dimensionless parameters with *T* = 1 and *γ* = 1, where *γ* is the friction coefficient, and a simulation timestep of *δt* = 0.002. To map simulation time to real time, we mapped one simulation timestep to 3.4 × 10^−6^ µs such that the self diffusion time of one bead in our model matched the self diffusion time of two attached amino acids (with size ≈ 0.76 nm) in water at room temperature (see Supplemental Material). At least 34 µs were used to equilibrate the simulations, where equilibration was verified by inspection of the radius of gyration.

### B. Density Functional Theory (DFT)

To investigate molecular interactions in a polymer system we have used classical density functional theory (DFT), a scheme based upon the minimisation of a dimensionless free energy functional ℱ that depends solely on the number density of beads *ρ*(*r*), and is written as ℱ[*ρ*(*r*)] [34]. In this work we have formulated DFT using mean-field theory so that the many-body polymer interactions are reduced to a single polymer interacting with a dimensionless mean field *w*(*r*). The optimum mean field minimizes ℱ and produces as output the equilibrium number density.

To model planar assemblies of FG nups, we formulated a 1D version of a previously successful 2D DFT formulation [22], where polymers were grafted onto the base of a cylinder with the assumption of rotational symmetry along the axial coordinate. This DFT has been previously described in extensive detail [15, 22, 28]. Here we describe the 1D version, consisting of polymers grafted onto a flat surface and assuming translational symmetry along this surface. The determining coordinate was therefore the height *z* above the grafting surface.

We took the approximation ℱ = ℱ_0_ + {ℱ_*vol*_ + ℱ_*att*_ + ℱ_*sur*_ + ℱ_*mf*_} where ℱ_0_ is the free energy functional describing a chain of *N* non-interacting point-like beads in the mean field, and excess terms representing excluded-volume, attractive, surface, and compensating mean field interactions respectively. ℱ_0_ is defined by the Hamiltonian

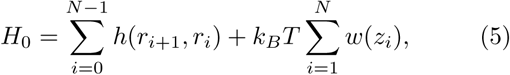

where *h* is a function (and *r* is the magnitude of bead separation) that imposes a rigid bond length of *r*_0_ between beads in a chain and *w*(*z*) is the 1D mean field.

ℱ_*vol*_ is the free energy functional imposing the excluded volume interactions between beads. To impose the excluded volume interactions, we used fundamental measure theory

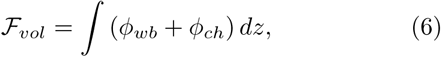

where *ϕ* _*wb*_ is the white bear functional [35] and *ϕ*_*ch*_ is the chain connectivity functional [36]. *ϕ* _*wb*_ is given by

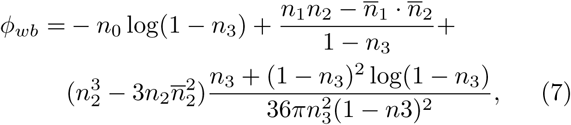

and *ϕ* _*ch*_ is

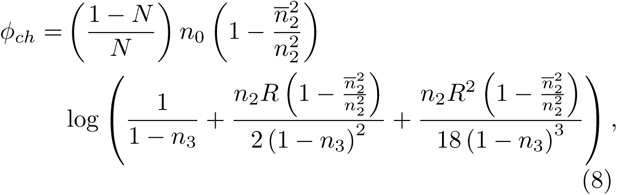

where *R* = *d/*2 is the bead radius and {*n*_*α*_} and 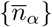 are sets of scalar and vector weighted densities, respectively, that are given by

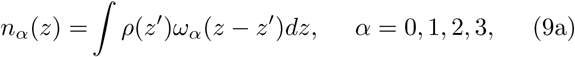

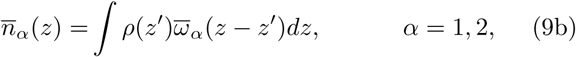

where *ρ*(*z*) is the one dimensional number density, *ω*_*α*_ and 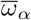 are the 1D geometrical weight functions of a sphere [37] given as 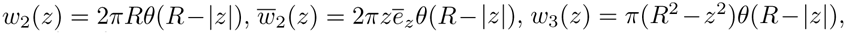 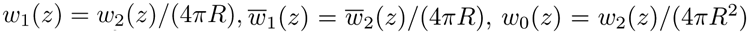, where *ē*_*z*_ is a unit vector and *θ* is the Heaviside function.

The cohesive term in the free energy, ℱ_*att*_, is implemented using the random phase approximation [38] and is given by

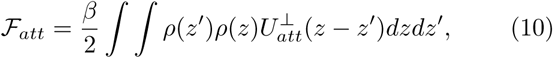

where *β* = 1*/k*_*B*_*T* and 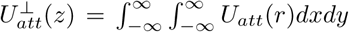 integrated over an infinite grafting area) with *r* being the magnitude of vector separation between beads, and *U*_*att*_ is given by equation 3.

The free energy term representing the interactions between beads and the surface is given as

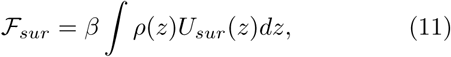

with *U*_*sur*_ as given in equation 1 in the Supplemental Material. The mean field energy, ℱ_*mf*_, is the dimensionless free energy term that compensates for the introduction of a mean field and is given as

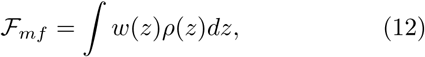

To incorporate a number of polymers, *N*_*p*_, one multiplies ℱ_0_ by *N*_*p*_ and interprets *ρ*(*z*) as the number density of *N*_*p*_*N* beads. To compute *ρ*(*z*), we solved the 1D diffusion equation for a random walk with contour length *Nr*_0_ in the presence of an external field *w*(*z*) [15]. We optimized *w*(*z*) through a discrete update rule

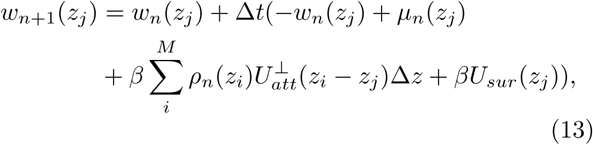

where *n* is an index representing the current iteration, Δ *t* is the update timestep, {*i, j*} are labels denoting discrete space, *µ* is the functional derivative of ℱ_*vol*_ with respect to *ρ*(*z*), and *M* is the total number of discrete spatial points (along the z axis). To ensure the stability of the update rule we used *M* = 1024 and Δ *z* = *z*_*max*_*/M* (*z*_*max*_ = 100 nm so that polymer beads were well within the spatial domain), Δ *t*= 0.001, and the initial mean field was set to zero for all *z*. Convergence was obtained when *w*_*n*+1_(*z*_*j*_) − *w*_*n*_(*z*_*j*_) ≤ 10^−7^ for all *j*.

## RESULTS

We model FG nups as freely jointed polymers consisting of *N* identical beads where one bead represents two amino acids (Figure 1a). FG nup excluded-volume (repulsive) and cohesive interactions are modelled through a short-ranged pair potential (equation 4) with a minimum value of *ϵ*_*pp*_, which is a measure of the cohesion strength between polymer beads. Thus defined, *ϵ*_*pp*_ is a phenomenological parameter capturing the general cohesive properties of FG nups arising from the different attractive interactions between the amino acids. We consider weak attractive interactions, with a *K*_*D*_ ∼ 0.1 M between two beads, as here found for 0.0 *k*_*B*_*T* ≤ *ϵ*_*pp*_ ≤ 1.0 *k*_*B*_*T* (see Figure S1b and Supplemental Material).

**FIG. 1.**
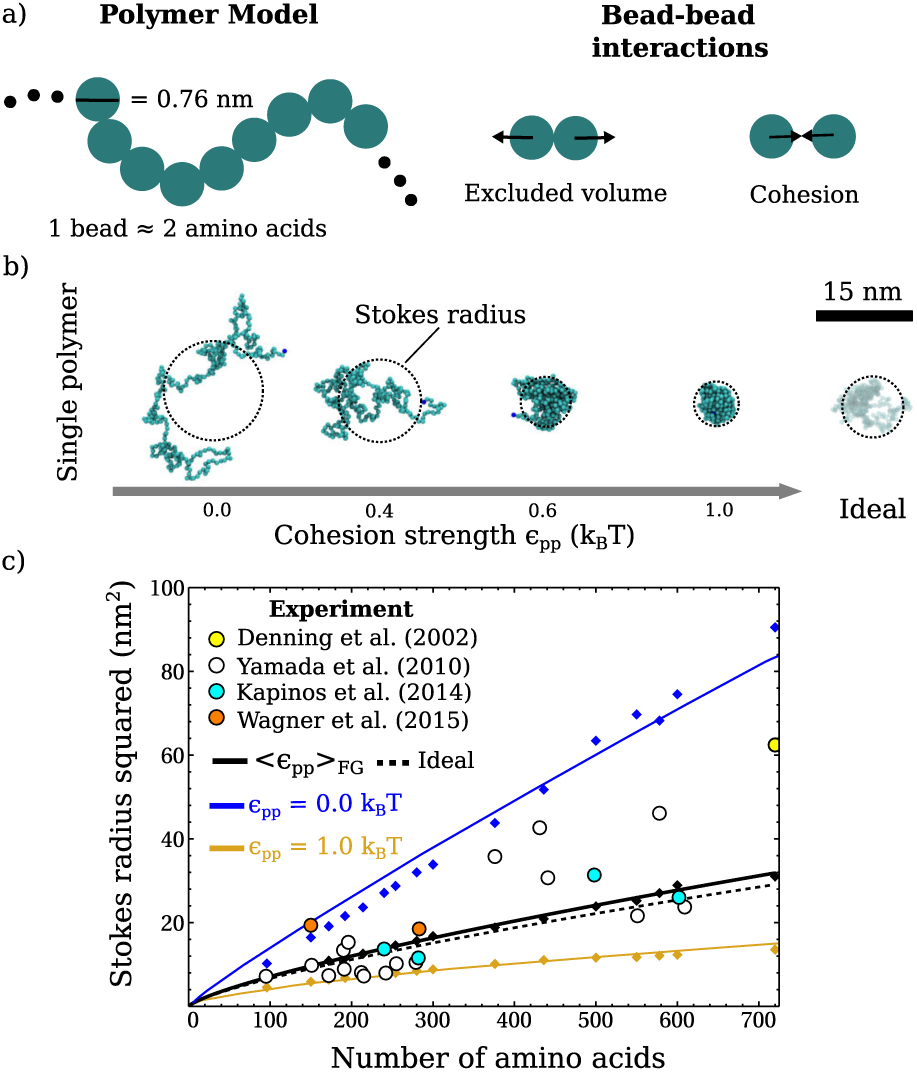
Parametrization of the polymer model by comparison with experimental Stokes radii [5, 10, 26, 27]. **a)** Illustration of the polymer model (MD). **b)** MD snapshots of a polymer (*N* = 300 beads) as a function of *ϵ*_*pp*_, alongside an ideal polymer. **c)** Experimental Stokes radii (circles) plotted against FG nup sequence lengths (see Table S1), compared with MD data (diamonds) alongside power law fits of the simulation data (solid and dashed lines).

Using an approach previously reported in the literature [39], the physiologically relevant *ϵ*_*pp*_ was determined by calculating hydrodynamic (Stokes) radii of specific FG constructs using MD simulations, and by comparing these with experimental data [5, 10, 26, 27] (see Table S1 for FG nups used and resulting *ϵ*_*pp*_ values).

Qualitatively, at *ϵ*_*pp*_ = 0.0 *k*_*B*_*T* (no cohesion) a polymer is in a swollen state; for 0.0 *< ϵ*_*pp*_ *<* 1.0 *k*_*B*_*T* polymers adopt morphologies that become increasingly compact with increasing *ϵ*_*pp*_; and at *ϵ*_*pp*_ = 1.0 *k*_*B*_*T* (“high cohesion”) a polymer forms a tight ball-like morphology (Figure 1b and Video S1). When all bead-bead interactions are nullified (ideal polymer), the polymer sizes lie – as expected – between the predictions for polymers with no and high cohesion (with excluded-volume).

The average of the parametrized *ϵ*_*pp*_ values (see Table S1) yields ⟨*ϵ*_*pp*_⟩_*FG*_ = 0.5 ± 0.2 *k*_*B*_*T* (mean ± standard deviation, see Figure 1c). Remarkably the experimental data points for polymers with cohesion strength ⟨*ϵ*_*pp*_⟩_*FG*_ coincide with the data from ideal polymer simulations (Figure 1c). This is a manifestation of polymer behaviour that typically occurs at Θ-point conditions [25]. Comparing the simulations for ⟨*ϵ*_*pp*_⟩_*FG*_ with those for ideal polymers, we also find similar scaling exponents for Stokes radii (Table S2) which are slightly less than the scaling exponents for *R*_*G*_, as is known for finite length polymers [41].

Importantly, we observe that the match between the mean FG nup behaviour, with ⟨ *ϵ*_*pp*_⟩_*F G*_, and ideal polymer behaviour is robust against the choice of model. Using a model (with a bead size of 0.57 nm) with lower estimates of the protein volume and persistence length of FG nups, we find a smaller ⟨*ϵ*_*pp*_⟩_*FG*_ = 0.26 ± 0.06 *k*_*B*_*T*, yet for this ⟨*ϵ*_*pp*_⟩_*FG*_ we still observe an excellent match with the ideal polymer prediction (Figure S2). Generally, to match the experimental data, the FG nup swelling due to excluded-volume interactions (Figure 1b) needs to be counteracted by intramolecular cohesion. As a consequence, overestimates of protein volume, and hence of excluded-volume, lead to larger *ϵ*_*pp*_ values in our comparisons with the experimental data. We note that FG nup domains with higher charged/polar content appear more extended [5], which in our analysis translates into smaller *ϵ*_*pp*_ values. Interestingly, this further extension can be entirely explained by the larger excluded volume of these specific domains, *i.e*., without needing to take into account explicit electrostatic interactions (Figure S3). This suggests that the attractive effects of various numbers of hydrophobic, charged, and hydrophilic amino acid regions, defining the heteropolymer nature of FG nups (Yamada et al. 2010) can be suitably captured in our homopolymer model with varying values of one interaction parameter.

We also tested against experimental data on FG nups that are grafted, at physiological densities, onto a planar surface (polymer film) [10, 22, 27]. MD simulations (Figure 2a and Video S2) for ⟨*ϵ*_*pp*_⟩_*FG*_ = 0.5 ± 0.2 *k*_*B*_*T* yield FG nup film thicknesses (Figure 2b, green shading) that are in agreement with the experimental data (Figure S4), with the exception of three data points (see Supplemental Material for an explanation). With these three exceptions, experimental and computational data for ⟨*ϵ*_*pp*_ ⟩_*FG*_ again show close agreement with the predictions for ideal polymers.

**FIG. 2.**
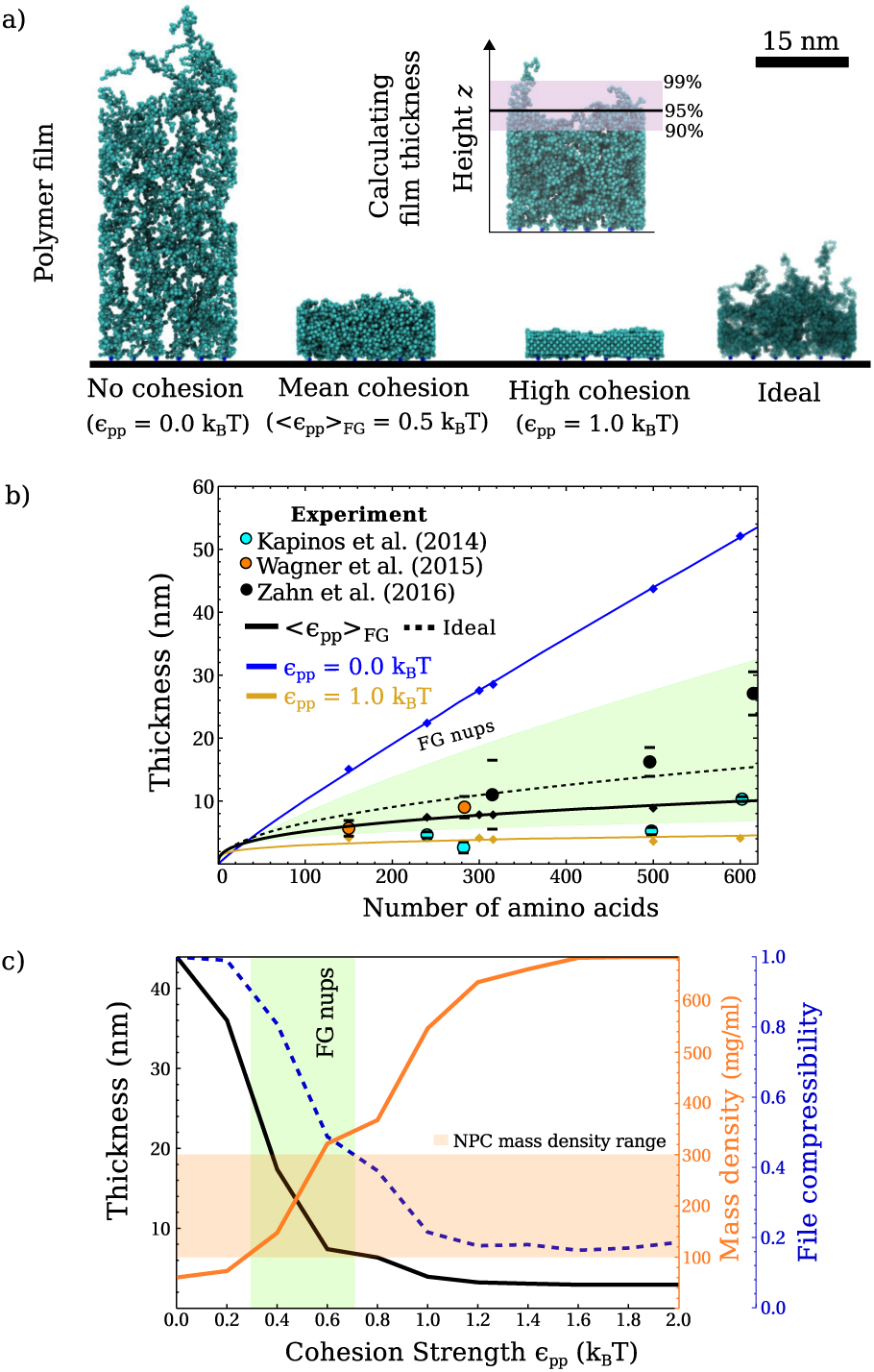
Comparing the parameterized polymer model with planar assemblies of FG nups. **a)** MD snapshots of polymers (*N* = 300 beads) grafted onto a surface (at a density of 3.2 polymers/100 nm^2^) for various interaction regimes. **b)** Experimental film thicknesses (circles) plotted against FG nup sequence lengths. MD simulation data (diamonds) are shown for *ϵ*_*pp*_ = {0.0, ⟨*ϵ*_*pp*_⟩_*FG*_, 1.0} *k*_*B*_*T* alongside ideal polymer data. **c)** Measures of compaction as a function of *ϵ*_*pp*_ (MD, see main text). The mass density range, from a simulated NPC, is shown in orange shading [40].

Next we explore how the identified ideal polymer morphology translates into functionally relevant properties in a polymer film assembly containing one type of FG nup (*N* = 300 beads). Firstly in the *ϵ*_*pp*_ range relevant for FG nups, we find that the computed film thickness (Figure 2c) shows a steep change from a swollen polymer film to a compact film. This is confirmed by other measures of film compaction, such as the mass density and file compressibility, where the file compressibility has been used to describe order in non-equilibrium and equilibrium many-body systems [42, 43]. The mass density is calculated by converting the number density within the film thickness to mg/ml assuming a molecular weight of 220 Da for a polymer bead, and the file compressibility corresponds to the losslessly compressed size (in Bytes) of a file containing the MD bead coordinates divided by the losslessly compressed size for *ϵ*_*pp*_ = 0.0 *k*_*B*_*T* (see Supplemental Material). The mass density of the FG nups, in the relevant *ϵ*_*pp*_ range, is consistent with the FG nup mass density in the NPC, as previously estimated using simulations which model the geometry of the nuclear pore scaffold, and the FG nups down to the amino acid level [40].

To further elucidate the molecular interactions within the polymer film assembly we use free energy density functional theory (DFT) modelling, in which grafted FG nups of the same type are described as ideal polymers interacting with a mean field that is optimized to best represent the net effect of excluded-volume, cohesion, and grafting of the FG nups to a surface [15]. This DFT yields film thicknesses that are in good agreement with the MD simulations of the same system (Figure S5), and allows us to quantify the effect of molecular interactions via the mean field per polymer in the film (Figure S6). Here the ideal polymer behaviour, where excluded-volume and cohesion balance out (Figure 3d), is articulated via the zero-crossings of the mean field energy per polymer and of the second virial coefficient in the relevant range of *ϵ*_*pp*_ (Figure 3).

**FIG. 3.**
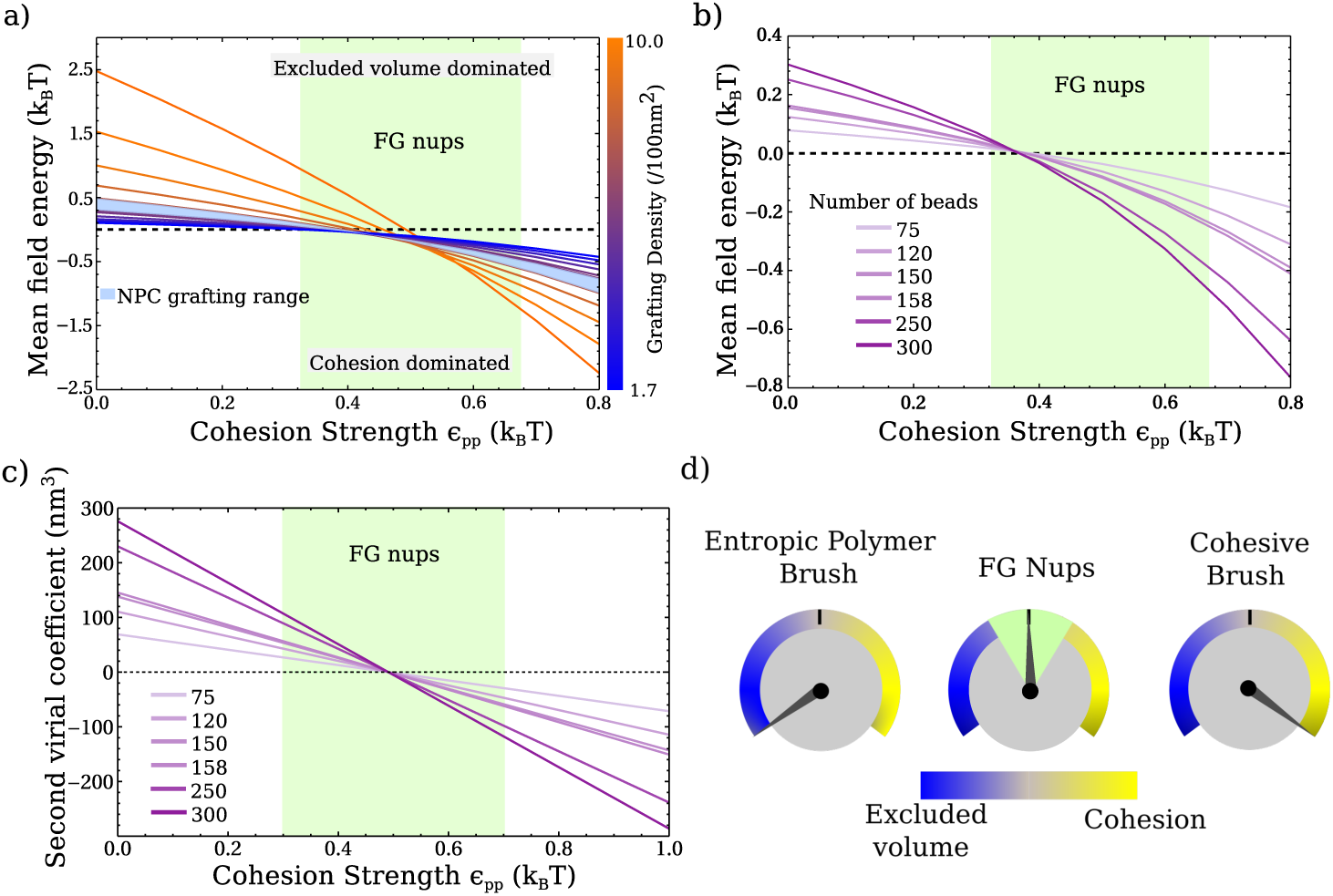
Planar assemblies of FG nups adopt morphologies that balance excluded-volume and cohesive interactions analysed by DFT. **a)** The mean field energy (see Equation 12, multiplied by *k*_*B*_*T*) divided by the number of polymers as a function of *ϵ*_*pp*_ for a range of grafting densities (*N* = 300 beads). **b)** Same as **(a)** but for various polymer lengths and for a physiological grafting density of 3.2 polymers per 100 nm^2^. **c)** The second virial coefficient per polymer (not per bead) as a function of *ϵ*_*pp*_ calculated from the bead-bead pair potential. **d)** An illustration of the balance between excluded-volume and cohesive interactions for FG nups.

We now investigate how the identified ideal polymer behaviour translates into a pore assembly, as here probed by MD simulations on NPC-mimetics based on well-defined numbers and types of FG nups that are grafted in a DNA origami pore scaffold, with an inner radius (∼20 nm) comparable to that of the NPC [44]. We observe that the change in polymer compaction is even more abrupt (in the relevant *ϵ*_*pp*_ range) for FG nups that are assembled in a pore geometry (Figures 4 and S7, and Video S3). This change in compaction results from the increased affinity between the polymers, which causes an increase in local concentration. This increased local concentration further enhances the probability of intra-/intermolecular interactions to come into play, such that attractive interactions can further compact the polymers, thus causing to a highly nonlinear response to changes in *ϵ*_*pp*_. As demonstrated here, this phenomenon is particularly articulated when the majority of the polymer become confined in the nanopore confinement. Hence in the physiologically relevant parameter range as determined above, we observe that FG nup assemblies in a nanopore can undergo major conformational changes for only minor changes in intermolecular interactions [15, 45].

**FIG. 4.**
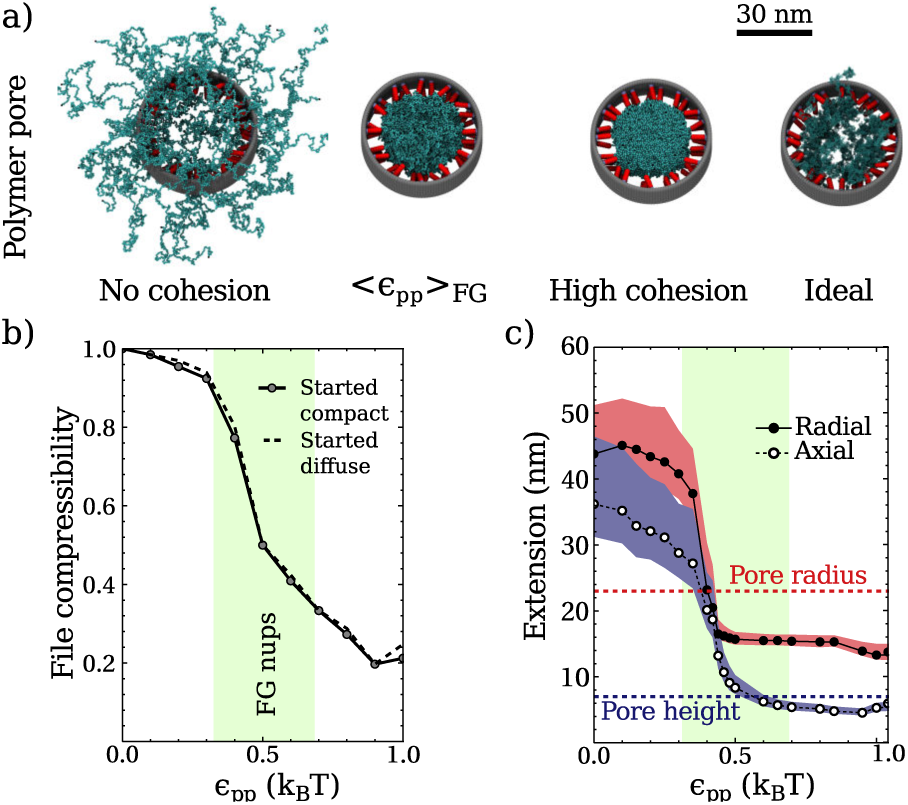
Investigating polymer morphology in an NPC-mimicking geometry containing one type of FG nup [44]. **a)** MD snapshots of polymers in a pore (polymers in blue; inner pore scaffold in grey; grafting handles in red) for various interaction regimes (48 polymers, *N* = 300 beads). **b)** File compressibility plotted as a function of *ϵ*_*pp*_. **c)** Polymer extension in the pore as a function of *ϵ*_*pp*_. The red (radial) and purple (axial) bands denote the extension thresholds that contain 99% and 90% of the total beads.

The polymer systems investigated thus far in this work have contained only one type of FG nup. However, it is known that the NPC contains many different types of FG nups with varying cohesiveness [5]. We therefore also investigated a binary polymer pore assembly, with 24 polymers with varying *ϵ*_*pp*_ and 24 non-cohesive polymers, and found that the presence of the non-cohesive polymers has a negligible effect on the morphology of the cohesive polymers (Figure 5).

**FIG. 5.**
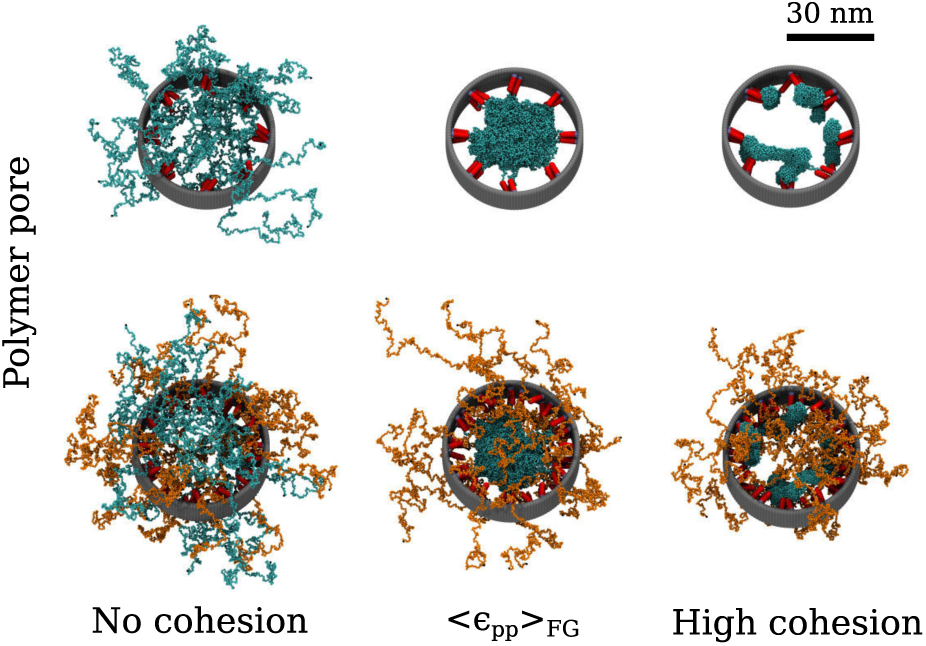
Incorporating polymers with no cohesion, but with excluded-volume interactions still present, into a polymer pore assembly does not change the morphology of the co-hesive polymers as compared with pores containing cohesive polymers only. (Top) MD snapshots of cohesive polymers (blue) (24 polymers, *N* = 300 beads). (Bottom) MD snap-shots of a binary mixture of 24 cohesive polymers (blue) and 24 non-cohesive polymers (orange) (*N* = 300 beads).

Thus far, we have observed that FG nup morphologies on average resemble those of ideal polymers, that they are at the transition between two (extreme) regimes of polymer behaviour, and that at this transition, it is possible to have large conformational changes for small changes in molecular interactions. This latter observation appears to be of significant physiological relevance, as it provides a mechanism by which FG nup assemblies in the NPC may open and close [15] to facilitate transport of large cargoes at millisecond time scales without compromising the transport barrier.

We next investigated the resealing dynamics of the barrier, *i.e*., how fast and by how much polymers fill the centre of the pore following a perturbation: holes (void of polymers) of 10, 20, and 30 nm diameter are created in an NPC-mimetic pore system containing 48 polymers [44] (Figures 6a and S8, and Video S4).

**FIG. 6.**
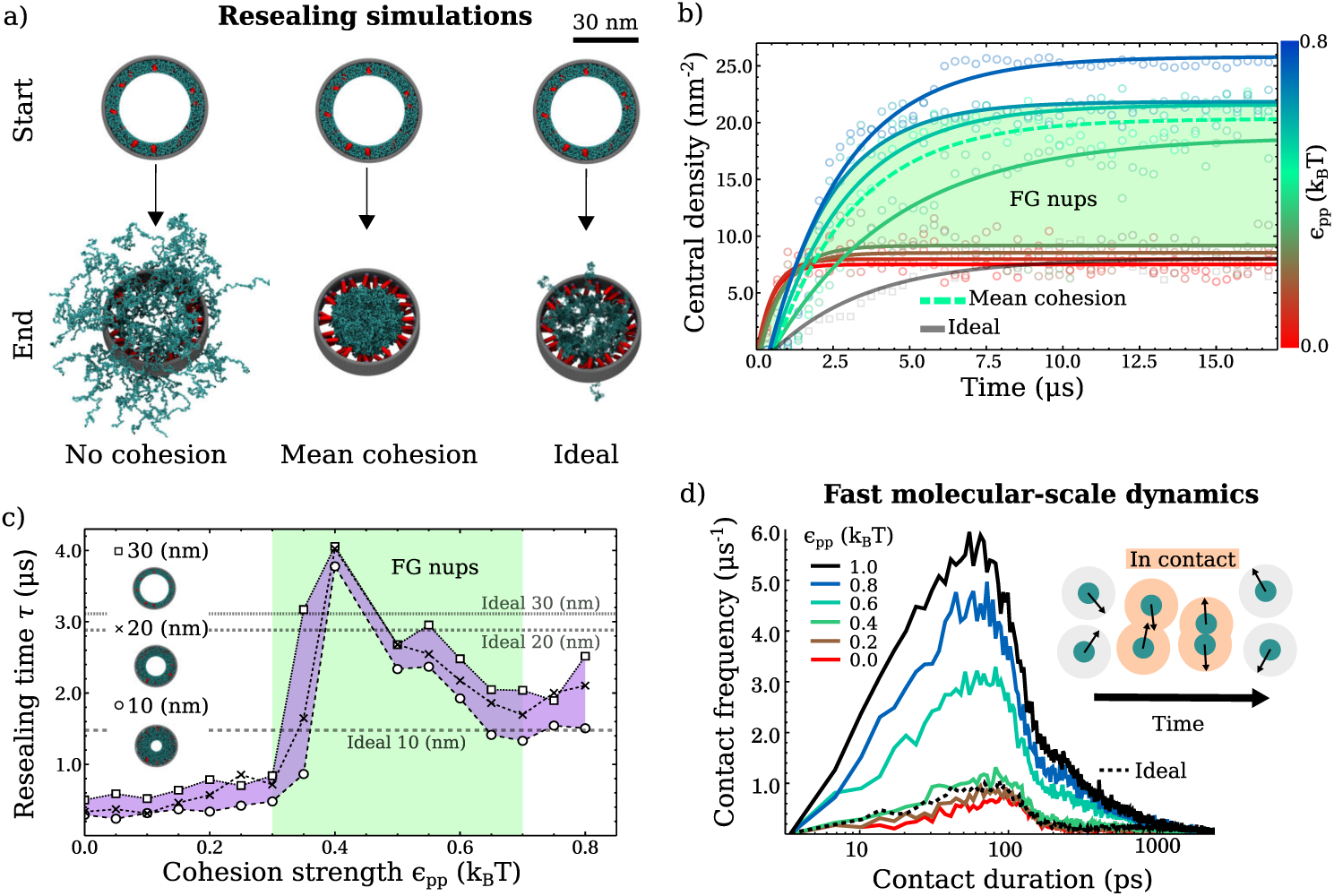
FG nup dynamics on the mesoscale and molecular scale. **a)** *In silico* polymer resealing in a nanopore [44]. The arrow represents 34 µs (or 10^7^ timesteps) of simulation time, pointing to resulting MD snapshots. **b)** The number of beads in a central circular cross-sectional area (5 nm radius) as a function of time, where the pore begins with a hole, 30 nm in diameter, at its centre. The cohesion strength colour bar represents the range *ϵ*_*pp*_ = 0.0 − 0.8 *k*_*B*_*T* (with 0.1 *k*_*B*_*T* increments). Lines represent fits using *ρ*(*t*) = *ρ*_*max*_(1 − exp(−*t/τ*)), where *ρ*_*max*_ is the maximum central density and *τ* is the resealing time (*N* = 300). **c)** *τ* as a function of *ϵ*_*pp*_ alongside resealing times for ideal polymers (dashed). **d)** Distribution of contact durations for a single isolated polymer (*N* = 300 beads).

We observe that FG nup resealing of the pore can be characterized by two regimes depending on the cohesion strength *ϵ*_*pp*_ (Figure 6b,c). For *ϵ*_*pp*_ ⪆ 0.4 *k*_*B*_*T*, the centre of the pore reaches a maximum polymer density that is approximately double that for *ϵ*_*pp*_ *<* 0.4 *k*_*B*_*T*. We also observe that for *ϵ*_*pp*_ ⪆ 0.4 *k*_*B*_*T*, the resealing rate appears slower than for *ϵ*_*pp*_ *<* 0.4 *k*_*B*_*T*, where polymers rapidly extend away from their tethering points (Figure S8). We probed the resealing time, *τ*, which is the time taken for the central density to reach ≈ 63 % its equlibrium value (Figure 6c). Notably, for polymers at ⟨*ϵ*_*pp*_⟩_*FG*_, the resealing time for “large cargoes” (exceeding 10 nm diameter [1–3]) is similar to that of ideal polymers, albeit that the central density remains at a relatively lower level for the ideal polymer case. This implies that the similarity to ideal polymers also extends to the dynamic behaviour of FG nups. Typically, the resealing times are on the microsecond time-scale, much faster – as needed to inhibit unspecific transport – than the millisecond time scale for transport events, and in agreement with previous simulations [46]. We also observe a sharp transition in the resealing time occurring at 0.3 *k*_*B*_*T < ϵ*_*pp*_ *<* 0.4 *k*_*B*_*T*, which falls in the relevant *ϵ*_*pp*_ range where we observed a drastic change in polymer structure (Figures 4b and c) and the ideal polymer behaviour (Figures 1c and 2b), while interesting it is not obvious what causes this observation.

Overall we can see that, in the physiologically relevant range, a small change in the cohesiveness can drastically change polymer morphology, which would contribute to the ease of the opening and closing of the pore. However, the time scale of the polymer dynamics is not greatly affected by changes in the cohesiveness (≤ 1.0 *k*_*B*_*T*), assuring fast resealing and rapid transportation of molecules.

Finally, we investigate how the cohesive interactions between FG nups affect dynamics at molecular and sub-molecular length scales. Nuclear magnetic resonance spectroscopy indicates fast (on the picosecond - nanosecond time scale) FG nup residue dynamics [47, 48], presumably indicative of entropically dominated FG nup behaviour. We simulated isolated polymers and measured the durations of bead-bead contacts, *i.e*., the elapsed time during which the centres of adjacent beads (excluding nearest bonded neighbours) are within the range of the attractive pair potential, (Figure 6d). Of note, the relevant time scales appear rather independent of the molecular interactions which is due to the weak (*ϵ*_*pp*_ ≤ 1.0 *k*_*B*_*T*) interaction strengths explored here. The strong dependence of the *collective* FG nup morphology on *ϵ*_*pp*_ (Figures 1, 2), is due to the rapid accumulation of many weak *individual* bead-bead contacts upon a moderate increase of *ϵ*_*pp*_, which can be attributed to the higher local bead concentration for larger *ϵ*_*pp*_. We observe that the overall dynamics does not depend on the exact shape of the model potential (Figure S9). Overall, individual FG nups exhibit fast dynamics (picosecond time-scale) at (sub-)molecular length scales, characteristic of their entropic nature. At larger length scales FG nup assemblies, dominated by the collective FG nup molecular interactions, exhibit microsecond dynamics with enhanced resealing (due to cohesion) which is of sufficient speed to maintain the transport barrier.

## CONCLUSION

In summary, we find that FG nup morphologies do not agree with predictions for purely entropic behaviour, dominated by (repulsive) excluded-volume interactions, nor with predictions for gel-like, strongly condensed behaviour governed by (attractive) cohesive interactions. This is fully consistent with previous studies [16, 21, 22, 24]. More surprisingly, we find that, on average, FG nup morphologies are consistent with the predictions for polymers that are characterized by the balance of repulsive and cohesive molecular interactions, *i.e*., the FG nups on average behave like ideal polymers, at or close to their Θ-point in physiological buffer solutions. FG nup mesoscale dynamics (microseconds) are also in agreement with predictions for ideal polymers, and are such that FG nup assemblies can reseal fast enough to reseal the barrier after transport events, which occur on millisecond time scales. Additionally, the rapid movement of FG nups on the molecular scale — maintained through a large range of cohesion strengths — will facilitate the uptake and release of nuclear transport factors and associated cargoes [49], whereas the accumulation of many weakly cohesive interactions facilitates the tight sealing of the NPC transport barrier. Taken together, this physical picture reconciles previous – apparently contradictory – identifications of FG nup assemblies as “entropic brushes” or “gels”, and provides a conceptual framework with which to interpret the transport selectivity of the NPC.

## ACKNOWLEDGMENTS

We thank Dino Osmanović (MIT), Roy Beck (Tel-Aviv), Larissa Kapinos (Basel), Roderick Lim (Basel), Ralf Richter (Leeds), and Anton Zilman (Toronto) for discussions. This work was funded by the Royal Society (A.Š.) and the UK Engineering and Physical Sciences Research Council (EP/L504889/1, B.W.H.).

