## Supplementary Information for "Intrinsically disordered nuclear pore proteins show ideal-polymer morphologies and dynamics"

### CONTENTS

Methods 1
Mapping self diffusion times 1
Single Polymer 1
Polymer film 1
Polymer pore 2
Calculating the dissociation constant 2
Model parameterization 2
Model validation 2
File compressibility 2
Calculating mass density 3
Resealing simulations 3
Calculating the second virial coefficient 3
WCA and Morse potential 3

Supplemental Notes 3
Film thickness discrepancies 3

Supplemental Tables, Figures, and Video legends 4

References 12

### METHODS

#### Mapping self diffusion times

To map the simulation timestep to a time scale relevant to real FG nups, we computed the diffusion coefficient, $D$, from simulations of 20 isolated beads (of radius $R = 0.38$ nm) and compared them with calculated diffusion constants for an alanine-proline dimer [1]. We fitted a straight line (with zero y-intercept) to the MSD which gave $D = (6.429 \pm 0.002\ \text{nm}^2)/6\Delta t$ where $\Delta t$ is the unknown unit of time. We obtained the unit of time, $\Delta t$, via the Einstein-Stokes relation: $D = \frac{k_B T}{6\pi\eta R}$, where $k_B$ is Boltzmann's constant, $T$ is the temperature, $\eta$ is the viscosity, and $R$ is the radius of the bead. Using $k_B T = 4$ pNnm and $\eta = 8.90\times10^{-10}$ pNs/nm$^2$ for water at room temperature implied that the unit of time of the simulations was $\Delta t = 1.707 \times 10^{-9} s$. This produced a translational diffusion constant of $D = 0.627 \times 10^{-9}$ m$^2$s$^{-1}$ for a bead which is comparable to diffusion constants calculated, from all-atom MD simulations with explicit solvent, for an alanine-proline dimer [1]. The elapsed simulation time is given by $t = N_s \times \delta t \times \Delta t$ where $\delta t = 0.002$ is the LAMMPS time step and $N_s$ is the number of iterations for which a configuration of the system is recorded.

*
†

#### Single Polymer

To simulate an isolated polymer in the solvent we placed the first bead at the origin of a simulation box with side lengths of 600 nm. For these simulations a polymer started in an extended conformation where beads formed a straight line.

#### Polymer film

The first bead of each polymer was fixed to a specific point, $\mathbf{r}_\perp = (x_\perp, y_\perp, z_\perp = 0.0\ \text{nm})$, on a triangular lattice in the $x-y$ plane, where $(x_\perp, y_\perp)$ represents a unique (for each polymer) vertex on the lattice. We approximated a polymer film with infinite extension in the $x$ and $y$ directions through periodic boundary conditions. The hardness of the surface is implemented through a bead-surface interaction given by

$$U_{sur}(\Delta r_\perp) = \begin{cases} 4\epsilon_{LJ}\left[\left(\frac{\sigma}{\Delta r_\perp}\right)^{12} - \left(\frac{\sigma}{\Delta r_\perp}\right)^{6}\right] + \epsilon_{LJ}, & \Delta r_\perp < d, \\ 0, & \Delta r_\perp \geq d, \end{cases} \tag{1}$$

where $\Delta r_\perp$ is the distance between a bead to the nearest point on the surface passing through the tethering points. Polymers were grafted onto the surface at a density of 3.3 polymers per 100 nm$^2$, which is in the range of the grafting density of a yeast NPC that is thought to be 3.1 to 4.1 polymers per 100 $nm^2$ (i.e. 5.2 - 6.9 pmol per $cm^2$) [2]. The surface area was set to 545 $nm^2$ and 18 polymers were used. The initial condition was generated using a short MD simulation where polymer beads were attracted to the surface.

#### Polymer pore

We based the pore geometry on an NPC-mimetic system (NuPOD) that we have used previously [3]. In this system, 48 polymers were grafted onto freely rotating rods that were fixed on the inside of a stationary cylinder. The rods and cylinder consisted of beads and all polymer, rod, and cylinder beads interacted through excluded-volume interactions only. In addition, the beads of the polymers interacted through an attractive potential as outlined above (equation 4 in main text).

#### Calculating the dissociation constant

We calculated the dissociation constant $K_D$ (in Molars) for two beads, in a simulation box with periodic boundary conditions, with cohesion strength $\epsilon_{pp}$. We used the relation $K_D = K_A^{-1} = (N_A n_1 (V - V_D)/n_0)^{-1}$ [4], where $K_A$ is the association constant, $N_A$ is Avogadro's number, $n_1$ is the number of simulation trajectories containing two beads that form a dimer, *i.e.*, the inter-particle separation is $\leq r_D$ where $r_D$ is the dimerization cut-off, $V$ is the volume of the simulation box ((6 nm)$^3$), $V_D = 4\pi r_D^3/3$ is the dimerization volume of one bead (with a volume, excluding the dimerization volume, per bead $(V - 2V_D)/2$ of 106-108 $nm^3$ for $r_D$ between 0.76-0.38 nm respectively [4]), and $n_0$ is the number of simulation trajectories containing two beads that have an inter-particle separation $> r_D$.

#### Model parameterization

To set the cohesion strength $\epsilon_{pp}$ for polymers with lengths corresponding to FG nups (Table S1) we performed MD simulations of single polymers with $\epsilon_{pp} = (0.0, 0.2, 0.4, 0.6, 0.8, 1.0)\ k_BT$. We computed the ensemble averaged Stokes radius $R_S$, the radius of a sphere that has the same diffusion coefficient as the polymer, from simulations run for 68 µs, using the HYDRO++ program (version 10) [5] assuming a temperature of 20.0 ℃, a solvent viscosity of 0.01 $gcm^{-1}s^{-1}$, and solution density of 1.0 g/$cm^3$. In order to find the $\epsilon_{pp}$ for a particular FG nup in this model, we interpolated the calculated Stokes radii as a function of $\epsilon_{pp}$ and solved for the $\epsilon_{pp}$ that yielded the experimental Stokes radius.

#### Model validation

To validate the parameterized polymer model we simulated polymer films at cohesion strengths $\epsilon_{pp}$ =( 0.0, 0.2, 0.4, 0.6, 0.8, 1.0) $k_BT$ and calculated the model film thickness predictions by numerically solving $\int_0^{z_\kappa} \rho(z)dz / \int_0^\infty \rho(z)dz = \kappa$, where $\kappa$ is the chosen fraction of beads and $z_\kappa$ is the distance above the surface that contains $\kappa N_p N$ beads. The film thickness is calculated with $\kappa = 0.95 \pm 0.05$, similar to previous comparisons between computational and experimental data [2]. The max height used for the density profile was 250 nm with 0.5 nm/bin for the histogramming. To obtain the film thickness at the NPC grafting density from the experimental data, we first fitted datasets, for a range of grafting densities, to a function $Z(D) = AD^B$, where $Z$ is the thickness, $D$ is the grafting distance between polymers, and $\{A, B\}$ are fitting parameters. We assumed that the average film thickness goes to zero for $D \gg 1$ nm. The experimental film thickness at the NPC grafting distance is then defined as $Z(D_H)$ where $D_H = \sqrt{\frac{1.15 \cdot 10^3}{6g}}$ is the grafting distance, $g = 5.4$ pmol/$cm^2$ is the approximate NPC grafting density, and the numerical factors convert $g$ to a grafting distance on a triangular lattice in units of nm.

#### File compressibility

To quantitatively compare polymer film and pore assemblies, we used a measure of bead compaction that could be implemented in the same way irrespective of geometry. We used the file compressibilty quantity which represents how much a data file containing bead coordinates can be losslessly compressed [6]. To compute the file compressibility we first calculated the minimum volume of a box, $V_0$, that contained all polymer beads in a system with $\epsilon_{pp} = 0.0\ k_BT$. $V_0$ was then discretized into cubes with side length $l = 0.76$ nm. For a simulation run at a certain $\epsilon_{pp}$, every 0.34 µ$s$ the simulation box was raster scanned to check if a cube was inside a bead (= 1) or not (= 0), and a file was generated containing all the states of the cubes. These files were then losslessly compressed using the bzip2 facility [7] and the average of the file sizes, $\varrho_{ave}(\epsilon_{pp})$, were computed. The file compressibility parameter was defined as $\varrho = (\varrho_{ave}(\epsilon_{pp}) - \varrho_{min})/(\varrho_{ave}(0.0) - \varrho_{min})$, where $\varrho_{min}$ is a reference compressed file size of a maximally ordered file containing a sequence of 0s only. For all cases the total number of characters in the uncompressed state file was $V_0/l^3$.

#### Calculating mass density

To compute the mass density, $\rho'$, in mg/ml in the polymer film assemblies we used the following formula

$$\rho' = \left( \frac{2\int_0^Z \rho(z)dz}{ZA^2} \right) B, \quad (2)$$

where $Z$ is the height (nm) above the grafting surface containing 95% of the polymer beads, $A$ is the area (nm$^2$) of the grafting surface, and $B = 110 \times 1.6605 \times 10^{-27} \times 10^{24} \times 10^3 = 182.655$ is a constant containing unit conversions and the average molecular weight of one amino acid (110 Da).

#### Resealing simulations

To investigate the parameterized model in a dynamic setting, we investigated the dynamic behaviour of polymer pore assemblies by performing 'resealing' simulations, where a hole (void of polymer beads) with diameter $d_{hole}$ is made at the centre of the polymer-coated pore and 'resealing' begins when this constraint is removed. The initial hole was created through a simulation where a cylinder with radius $d_{hole}/2$ and length $L = 200$ nm was dragged through the polymers inside the pore. The cylinder was a continuum, *i.e.*, not constructed of point like beads, and the distance between a bead and the cylinder was defined as the distance from the bead to the nearest point on the surface, therefore the force the surface exerted on the bead was along the direction that is normal to the surface at that point. The cylinder was placed at an initial height so as not to make contact with any bead, and with its axis permanently aligned with the pore scaffold. The centre of the cylinder was then incrementally moved towards the origin. The cylinder and beads interacted through excluded-volume only using the WCA pair potential given by equation 1 in the main text. Whilst the cylinder was moving an external constraint was imposed to force polymer beads to within the axial dimensions of the pore scaffold, without causing overlapping between beads. The simulation imposing the constraints was run for 0.34 µs and then a second simulation, marking the beginning of resealing, was performed with the artificial cylinder and external constraints removed. The resealing of the pore is quantified through a central density, *i.e.*, the number of beads in a circle of radius 5 nm located at the origin, which we compute every 0.34 µ$s$, and we fit this data to a relaxation function $\rho(t) = \rho_0 (1 - \exp(-t/\tau))$, where $\rho$ is the central density, $\rho_0$ the equilibrium central density, $t$ is time, and $\tau$ is the resealing time: the time taken for the central density to reach $\approx 0.63\rho_0$.

#### Calculating the second virial coefficient

We considered two beads, treated as weakly attractive ($\epsilon_{pp} \leq k_B T$) hard spheres, interacting with a pair potential given by equation 4 in the main text. The second virial coefficient per bead, $B_2(\epsilon_{pp})$, is given as

$$\begin{aligned} B_2(\epsilon_{pp}) &= 2\pi \left( b + \int_{r_0}^{r_c} r^2 \left(1 - \exp\left(-U_{att}(r)/k_B T\right)\right) dr \right), \\ &\approx 2\pi \left( b + \int_{r_0}^{r_c} r^2 \frac{U_{att}(r)}{k_B T} dr \right), \end{aligned} \quad (3)$$

where $b = r_0^3/3$, $r_0 = d$ is the contact distance between two beads, and $r_c$ is the cut-off of the attractive potential ($= \infty$ for infinite ranged potentials) [8]. The second virial coefficient per polymer is calculated as $NB_2(\epsilon_{pp})$, where $N$ is the number of beads in one polymer.

#### WCA and Morse potential

To test whether the static and dynamic properties of polymer assemblies depend on the choice of pair potential, we repeated a selection of simulations with a different potential to that as described in the main methods. This potential consists of a WCA potential (excluded-volume) given in equation 1 in the main methods and a truncated and shifted Morse potential (attraction) given by

$$\begin{aligned} \phi(r) = \epsilon'_{pp} &\exp(\alpha(-2(r + r_c) + r_0))(-2\exp(\alpha(2r_c + r)) +, \\ &\exp(\alpha(r_0 + 2r_c)) + \exp(\alpha(2r + r_0))(2r\alpha - 1 - 2\alpha r_c), \\ &+ \exp(\alpha(2r + r_c))(2 - 2r\alpha + 2\alpha r_c)) \end{aligned} \quad (4)$$

where $\epsilon'_{pp} = 1.13024 \times \epsilon_{pp}$ corrects the minimum of the potential due to the truncation and shifting procedure, $\alpha = 6.0$ nm$^{-1}$, $r_0 = 0.76$ nm, and $r_c = 1.52$ nm. The resulting pair potential (see Figure S9a) is then given by

$$U(r) = \begin{cases} U_{vol}(r) + \phi(r), & r \leq r_0, \\ \phi(r), & r_0 \leq r \leq r_c. \end{cases} \quad (5)$$

### SUPPLEMENTAL NOTES

#### Film thickness discrepancies

MD film thicknesses as predicted from polymers with the parameterized cohesion strength $\langle \epsilon_{pp} \rangle_{FG} = 0.5 \pm 0.2\ k_B T$ (Figure 2b, green shading), are in agreement with the experimental data (Figure S4) with the exception of three data points. One data point (Nup214, blue symbol, 282 amino acids) has a film thickness that is less than the experimental Stokes radius and another data point (Nup62, blue symbol, 240) has a resulting $\epsilon_{pp}$ that is almost double its $\epsilon_{pp}$ from the single-molecule parameterization, putting into question the assumption of lateral homogeneity that underpins the experimental analysis for those source data [9], and the third anomalous data point is for non-glycosylated Nup98 (blue symbol, $\approx 500$ amino acids) [9] for which the – physiologically relevant – glycosylated version (Nup98-Glyco, black symbol, $\approx 500$ amino acids) yields a three times higher film thickness [2].

### SUPPLEMENTAL TABLES, FIGURES, AND VIDEO LEGENDS

| FG nup | No. of amino acids | Protein volume (nm$^3$) [a] | Polymer volume (nm$^3$) [b] | Charged /hydrophobic ratio [c] | Single polymer | | Polymer film | | Source |
|---|---|---|---|---|---|---|---|---|---|
| | | | | | $R_S$ (nm) | $\epsilon_{pp}$ (kT) | Thickness (nm) | $\epsilon_{pp}$ (kT) | |
| Nsp1 | 95 | 9.1 | 10.8 | 1.31 | 2.68 | 0.50 | - | - | [11] |
| Nsp1p-5FF | 150 | 15.3 | 17.0 | 1.27 | 4.4 | - [d] | 5.68 | 0.54 | [12] |
| Nup60 | 151 | 15.3 | 17.2 | 0.95 | 3.13 | 0.47 | - | - | [11] |
| Nsp1n | 172 | 15.2 | 19.7 | 0.08 | 2.71 | 0.69 | - | - | [11] |
| Nup100s | 190 | 19.3 | 21.72 | 1.0 | 3.66 | 0.42 | - | - | [11] |
| Nup145Ns | 191 | 19.4 | 21.8 | 0.89 | 2.98 | 0.60 | - | - | [11] |
| Nup116s | 196 | 20.7 | 22.4 | 1.35 | 3.91 | 0.36 | - | - | [11] |
| Nup42 | 212 | 18.5 | 24.3 | 0.14 | 2.84 | 0.72 | - | - | [11] |
| Nup49 | 215 | 18.4 | 24.6 | 0.13 | 2.69 | 0.93 | - | - | [11] |
| Nup62 | 240 | 20.5 | 27.5 | 0.03 | 3.7 | 0.49 | 4.62 | 0.91 | [9] |
| Nup145N | 242 | 21.7 | 27.7 | 0.14 | 2.82 | 0.85 | - | - | [11] |
| Nup57 | 255 | 22.3 | 29.2 | 0.14 | 3.19 | 0.61 | - | - | [11] |
| Nup1c | 279 | 24.8 | 31.9 | 0.14 | 3.24 | 0.63 | - | - | [11] |
| Nup214 | 282 | 23.8 | 32.3 | 0.14 | 3.4 | 0.60 | 2.64 | 1.10 | [9] |
| Nsp1p-12FF | 283 | 27.9 | 32.3 | 1.4 | 4.3 | 0.42 | 9.03 | 0.47 | [12] |
| Reg-FSFG | 315 | 27.8 | 36.4 | 0.0 | - | - | 11.01 | 0.44 | [2] |
| Nup2 | 376 | 36.7 | 43.0 | 1.1 | 5.98 | 0.15 | - | - | [11] |
| Nsp1m | 431 | 40.9 | 49.4 | 1.22 | 6.53 | 0.19 | - | - | [11] |
| Nup159 | 441 | 40.6 | 50.6 | 0.69 | 5.54 | 0.40 | - | - | [11] |
| Nup98-Glyco | 496 | 45.1 | 56.9 | 0.2 | - | - | 16.22 | 0.41 | [2] |
| Nup98 | 498 | 44.7 | 57.1 | 0.2 | 5.6 | 0.40 | 5.24 | 0.72 | [9] |
| Nup116m | 551 | 48.9 | 63.2 | 0.11 | 4.65 | 0.50 | - | - | [11] |
| Nup1m | 578 | 57.0 | 66.3 | 1.08 | 6.79 | 0.38 | - | - | [11] |
| Nup153 | 602 | 55.7 | 69.1 | 0.5 | 5.1 | 0.48 | 10.31 | 0.50 | [9] |
| Nup100n | 609 | 54.9 | 69.9 | 0.11 | 4.87 | 0.49 | - | - | [11] |
| Nsp1 | 615 | 57.4 | 70.4 | 0.93 | - | - | 27.09 | 0.34 | [2] |
| Nup2p | 720 | 72.1 | 82.6 | 1.11 | 7.9 | 0.28 | - | - | [13] |

[a] Sum of Van der Waals volumes for each specific amino acid [10].
[b] Sum of bead volumes. A diameter of 0.76 nm is used for all calculations.
[c] Based on the same amino acid classification as used in [11].
[d] Experimental Stokes radius is slightly overestimated due to polydispersity [12].

TABLE S1. Characterisation, and experimental dimensions (Stokes radius $R_S$ and film thickness), of the resulting cohesion strengths ($\epsilon_{pp}$, from MD) for all FG nups used in this work.

| Interaction regime | $R_S = A_0(N_{AA})^\nu$ | | $R_G = A_0(N_{AA})^\nu$ | |
|---|---|---|---|---|
| | $A_0$ (nm) | $\nu$ | $A_0$ (nm) | $\nu$ |
| $\epsilon_{pp} = 0.0\ k_BT$ | $0.26 \pm 0.02$ | $0.55 \pm 0.01$ | $0.22 \pm 0.05$ | $0.61 \pm 0.03$ |
| $\epsilon_{pp} = \langle \epsilon_{pp} \rangle_{FG}$ | $0.47 \pm 0.04$ | $0.38 \pm 0.02$ | $0.4 \pm 0.1$ | $0.38 \pm 0.04$ |
| $\epsilon_{pp} = 1.0\ k_BT$ | $0.63 \pm 0.05$ | $0.27 \pm 0.01$ | $0.54 \pm 0.05$ | $0.24 \pm 0.02$ |
| Ideal | $0.35 \pm 0.03$ | $0.42 \pm 0.01$ | $0.25 \pm 0.04$ | $0.48 \pm 0.03$ |
| Experiment | $0.2 \pm 0.3$ | $0.5 \pm 0.2$ | - | - |

TABLE S2. Fitting parameters $\{A_0, \nu\}$ for $R_S$ and $R_G$ using a power law fit to MD simulation data for different values of $\epsilon_{pp}$ and for ideal polymers, as well as to the experimental data. Here, $N_{AA}$ is the number of amino acids. Note that the obtained values of the scaling exponent $\nu$ do depend on the choice of $A_0$ (here left as a free fitting parameter). The uncertainties represent 95% confidence intervals.

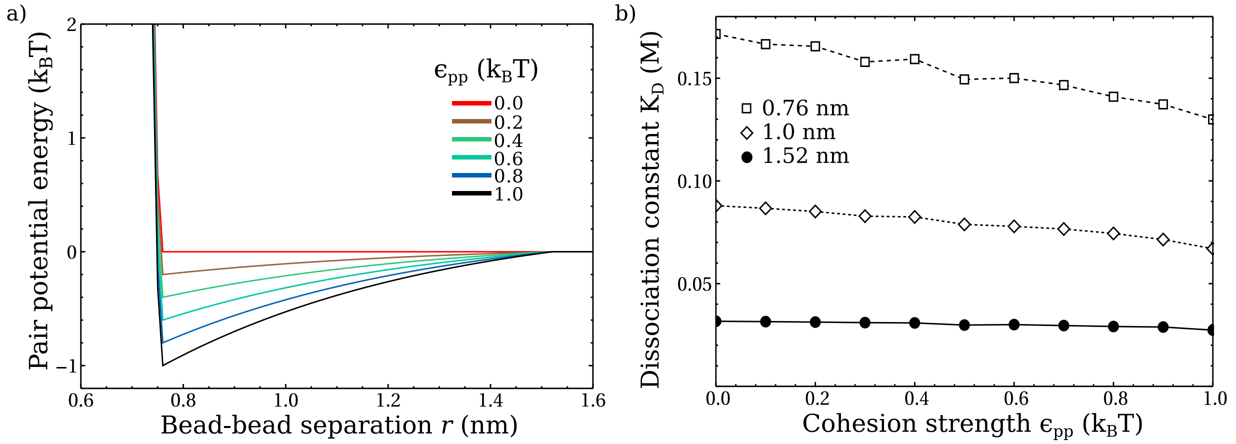


FIG. S1. Defining and quantifying the interactions between polymer beads. **a)** Plot of the total pair potential energy, $U_{pp}(r)$, between two polymer beads, with a diameter of 0.76 nm, at a centre-to-centre distance $r$, shown for various cohesion strengths $\epsilon_{pp}$. **b)** The dissociation constant $K_D$ between two individual polymer beads as a function of $\epsilon_{pp}$, for three choices of the dimerization cut-off (see Methods): the maximum inter-particle separation at which two beads are considered a dimer. For all choices of the dimerization cut-off, the number of simulation trajectories containing dimers was $> 2000$.

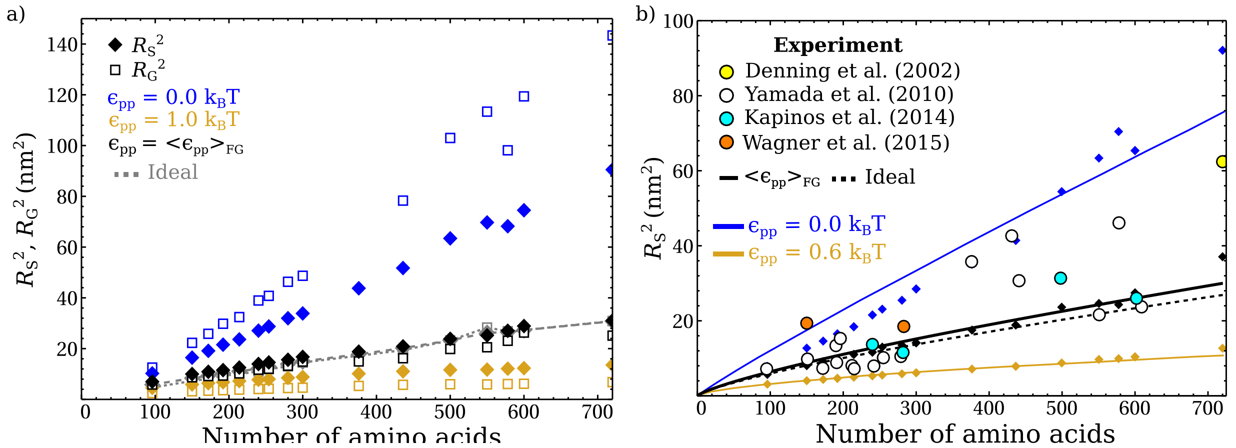


FIG. S2. Further quantification of single-molecule morphologies. **a)** Comparing the Stokes radii $R_S$ to the radius of gyration $R_G$ for various interaction regimes as calculated from MD simulations of polymers with a bead diameter of 0.76 nm. **b)** Comparing Stokes radii from MD simulations to experiments, as in Figure 1c but here using a bead diameter of 0.57 nm, *i.e.*, a polymer with a predicted persistence length smaller than that for FG nups (0.29 nm) and with an excluded volume that underestimates that of FG nups by $\approx 30\%$. The beads interacted through an attractive pair potential, as given in equation 3 (see main text), with $d$=0.57 nm and with a cut-off range $r_c = 1.52$ nm. In this model $\langle\epsilon_{pp}\rangle_{FG} = 0.21 \pm 0.06\ k_BT$. As for the model discussed in the main text, the behaviour for this $\langle\epsilon_{pp}\rangle_{FG}$ closely matched the predictions for ideal polymers.

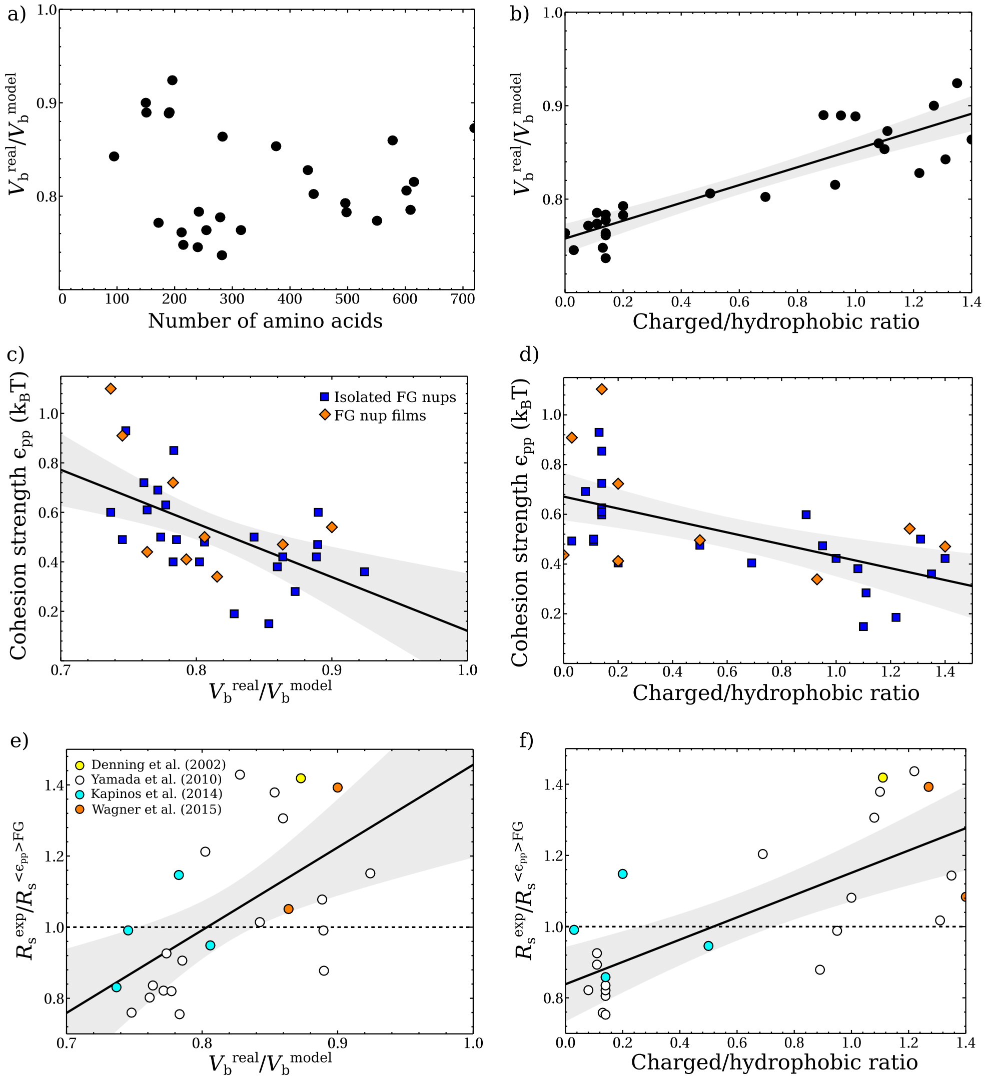


FIG. S3. Effects of excluded volume and ratio of charged/hydrophobic residues in the model. a) With our choice of parameters, the excluded volume is slightly overestimated, as here quantified by comparing the total sum of van der Waals volumes of the amino acids ($V_b^{\mathrm{real}}$) with the total excluded volume in our model ($V_b^{\mathrm{model}}$). Results are plotted for all analyzed sequences as a function of their sequence length. b) This $V_b^{\mathrm{real}}/V_b^{\mathrm{model}}$ is here shown to correlate with the ratio of charged to hydrophobic amino acids in these sequences, classified in line with previous work [11]. c) When $V_b^{\mathrm{real}}$ is larger compared with $V_b^{\mathrm{model}}$, a smaller $\epsilon_{pp}$ is found, as the experimental morphology of those FG nups is more extended. d) A similar dependence is found as a function of the charged/hydrophobic ratio of the FG nups. e,f) Ratio of the experimental Stokes radii and the Stokes radii predicted for $\langle\epsilon_{pp}\rangle$, as a function of $V_b^{\mathrm{real}}/V_b^{\mathrm{model}}$ (e) and of the charged/hydrophobic ratio (f). These data suggest that variations between FG nups can at least in part be attributed due to differences in the excluded volumes of their amino acids, in addition to or instead of differences in charge/hydrophobicity, as proposed previously [11]. The solid lines are linear fits, and the grey shadings are the corresponding 95% mean prediction bands.

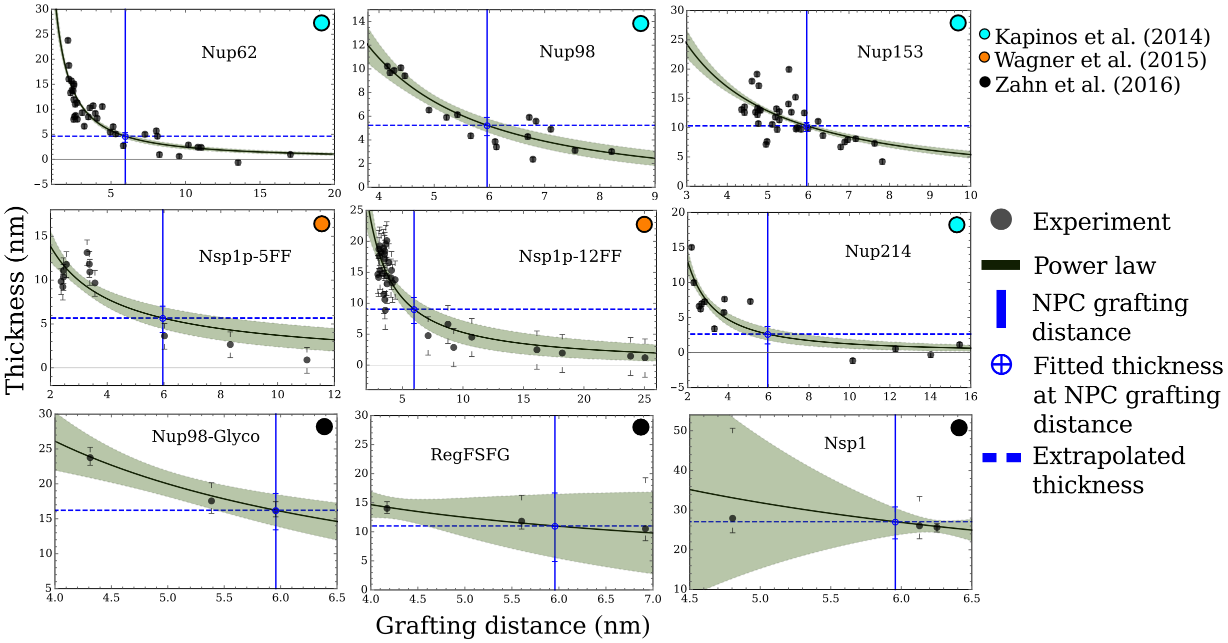


FIG. S4. Interpolation of the experimental data from FG nup films, to estimate the film thickness for a grafting distance/density that corresponds to the NPC (3.2 polymers/100 $nm^2$, or 5.4 pmol/$cm^2$). Data were fitted with a power law, with 95% mean prediction bands (green shading).

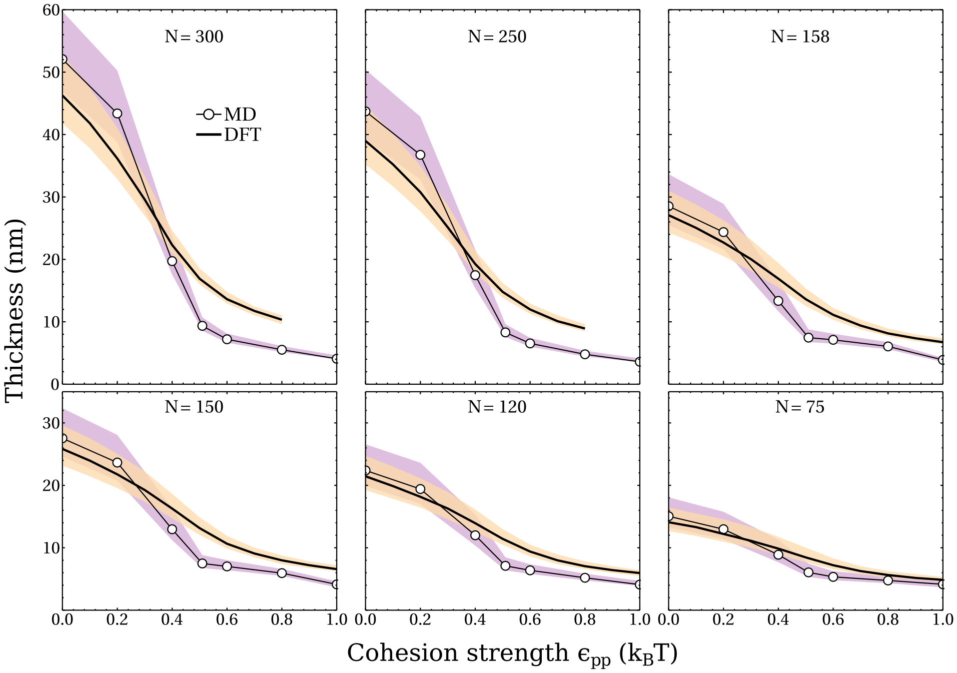


FIG. S5. Benchmarking DFT against MD film data for the thickness of FG nup films, as a function of cohesion strength $\epsilon_{pp}$. The bands denote a tolerance of ±5% of the total number of beads. DFT thicknesses at $\epsilon_{pp} > 0.8\ k_BT$ were unavailable for $N = 250$ and 300 beads, since the dense packing complicated the convergence of the DFT calculations.

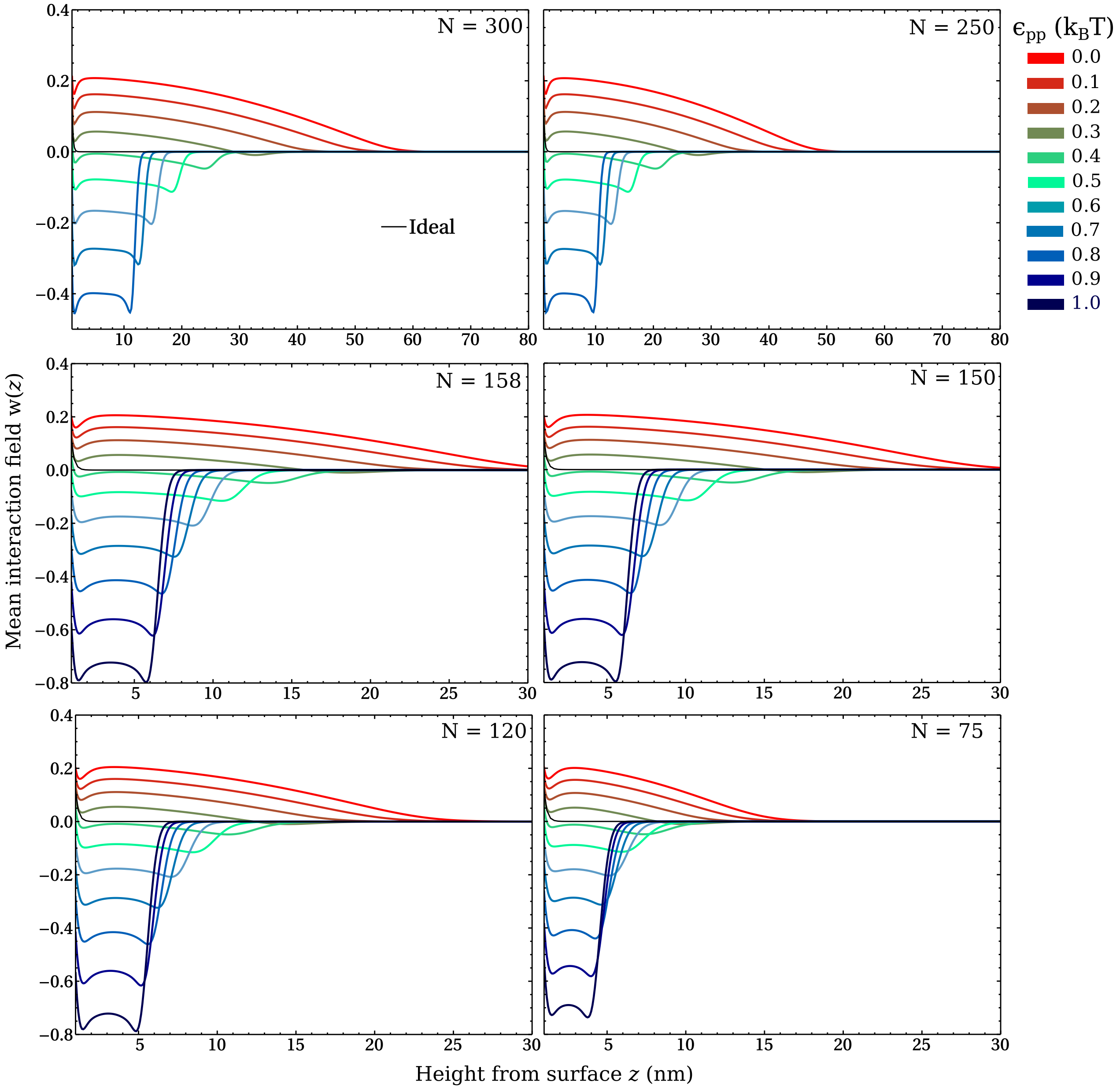


FIG. S6. DFT equilibrium mean fields $w(z)$ (see equation 5 main text) for polymer films comprising polymers at various chain lengths $N$ and cohesion strengths $\epsilon_{pp}$. We show the mean fields just above the grafting plane. The mean field energy per polymer follows by the integration (over $z$) of $w(z)\rho(z)$, divided by the number of polymers $N_p$ and multiplied by $k_B T$ (see equation 12 main text).

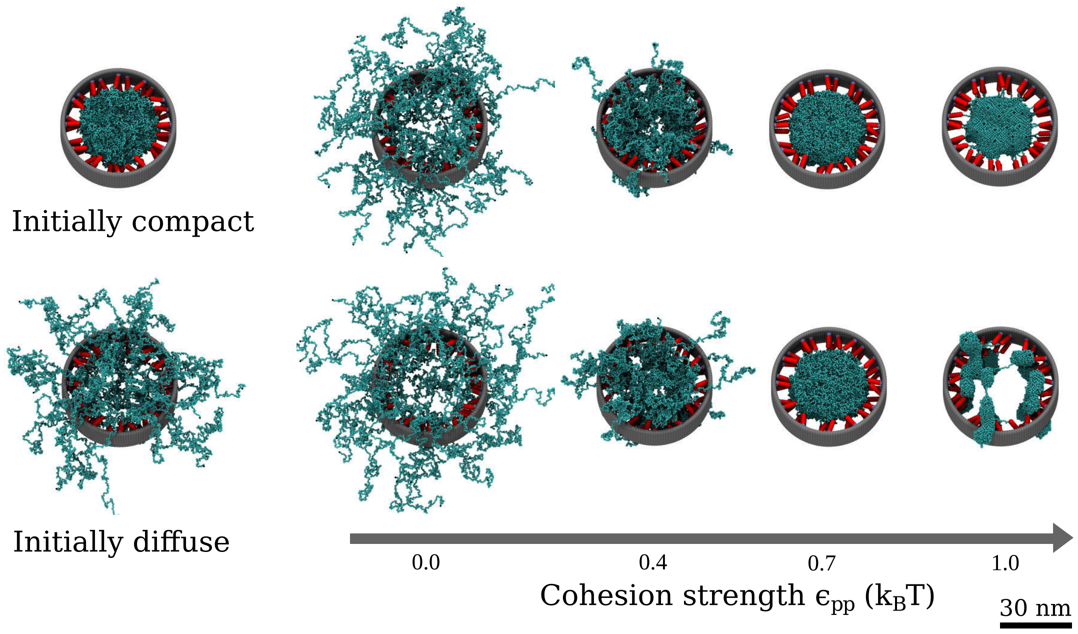


FIG. S7. Snapshots of polymer pore configurations (see Figure 4) for various $\epsilon_{pp}$ from two different initial conditions.

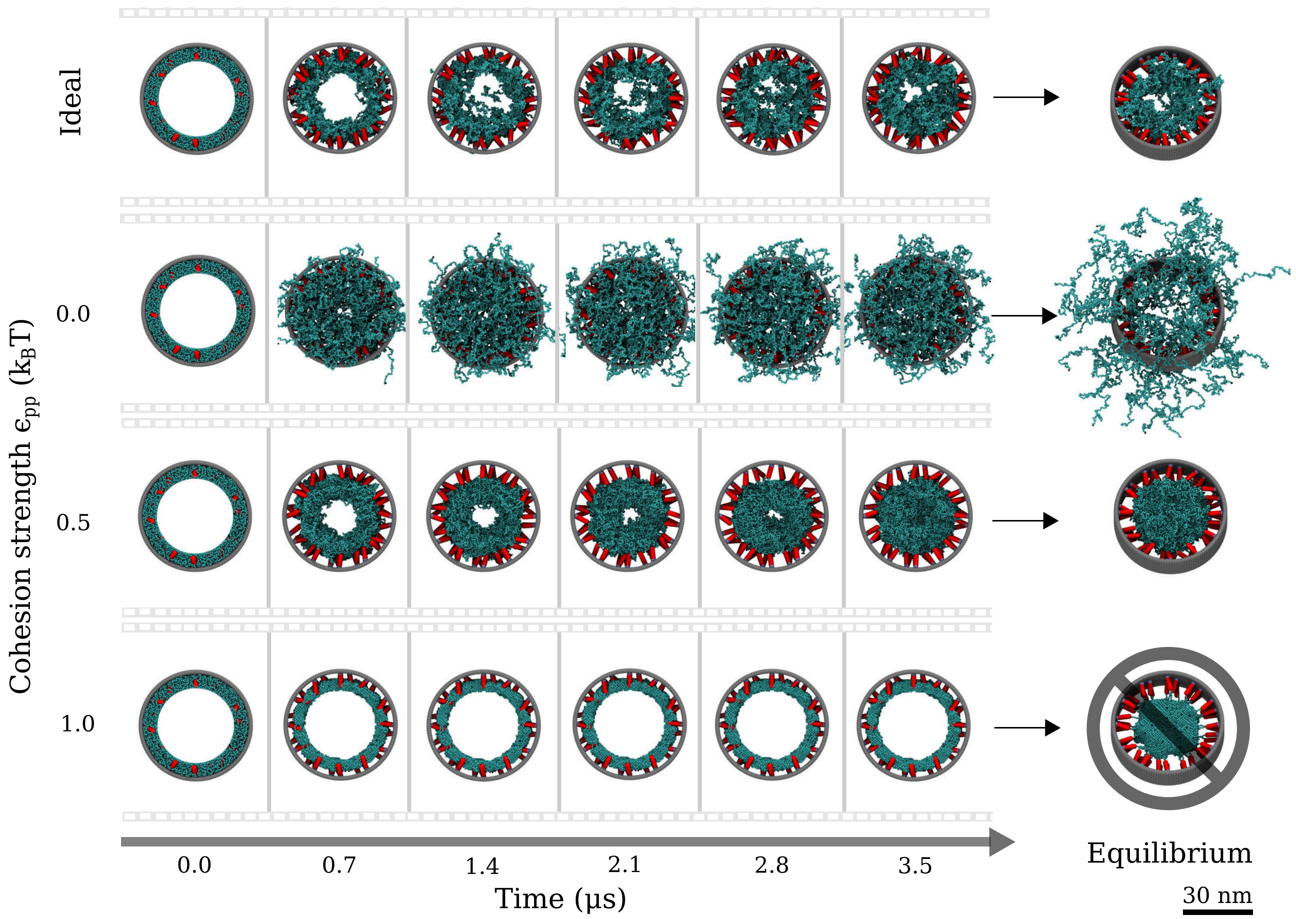


FIG. S8. Snapshots of polymer pore configurations (see Figures 4 and S7) from the dynamic resealing simulations (see Methods) and their resulting configurations after 34 μs of simulation time. For $\epsilon_{pp} = 1.0\ k_BT$ the pore did not converge to the configuration that was obtained by starting from a compact initial condition, hence the pore did not reseal.

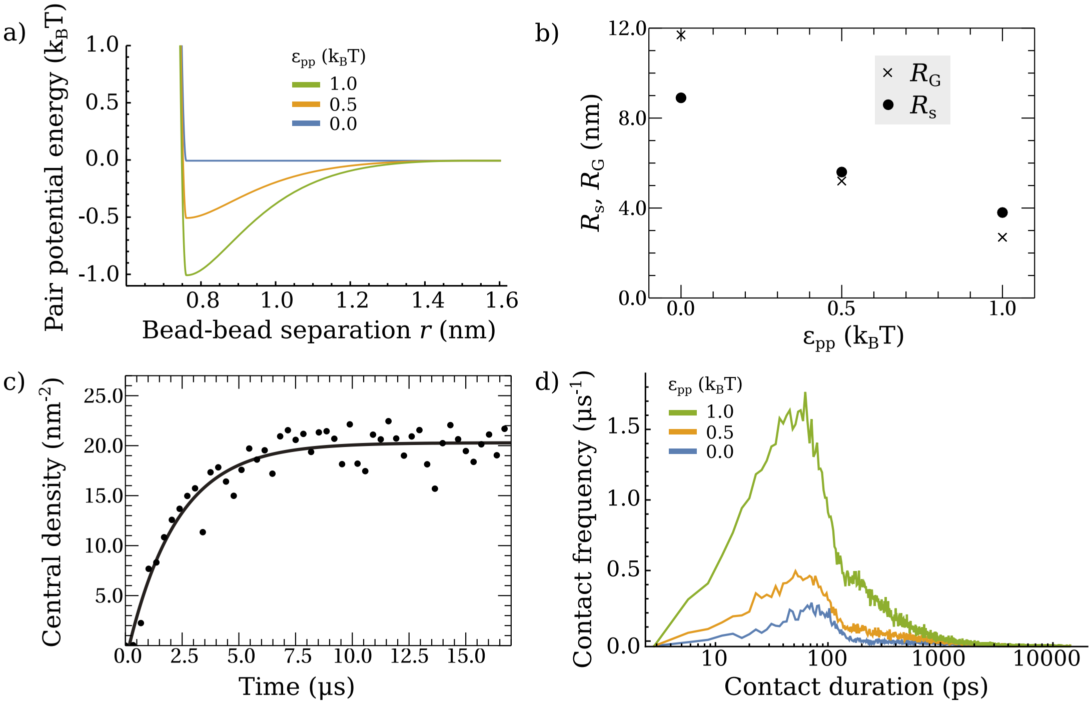


FIG. S9. Static and dynamic properties of polymers remain independent of the choice of the model potential, as shown here for a combined WCA (excluded-volume) and Morse (cohesion) potential. **a)** Pair potential energy as a function of bead-bead separation for three values of the cohesion strength $\epsilon_{pp}$. **b)** Hydrodynamic (Stokes) radius, $R_s$, and radius of gyration $R_G$ for $\epsilon_{pp} = (0.0, \langle\epsilon_{pp}\rangle_{FG} = 0.5, 1.0)\ k_BT$ ($N = 300$ beads). **c)** As in figure 6, time-dependence of the central areal density (5 nm radius from the centre of the pore) for a polymer pore assembly starting with a hole (30 nm in diameter) at the centre. The case for the cohesion strength that best matches experimental data, $\epsilon_{pp} = \langle\epsilon_{pp}\rangle_{FG}$, is shown. **d)** Distribution of contact durations for a single isolated polymer ($N = 300$).

**VID. S1.** Sequences of subsequent snapshots of single polymer MD simulations as shown in Figure 1. The order corresponds to a polymer with $\epsilon_{pp} = (0.0,\ 0.4,\ 0.6,\ 1.0)\ k_B T$ and an ideal polymer (with no excluded-volume and no cohesion).

**VID. S2.** Sequences of subsequent snapshots of polymer film MD simulations as shown in Figure 2. The order corresponds to polymers with $\epsilon_{pp} = (0.0,\ 0.5,\ 1.0)\ k_B T$ and ideal polymers (with no excluded-volume and no cohesion).

**VID. S3.** Sequences of subsequent snapshots of polymer pore MD simulations as shown in Figure 4. The order corresponds to pores containing polymers with $\epsilon_{pp} = (0.0,\ 0.5,\ 1.0)\ k_B T$ and ideal polymers (with no excluded-volume and no cohesion).

**VID. S4.** Sequences of subsequent snapshots of polymer pore resealing MD simulations as shown in Figure 6. The order corresponds to pores containing polymers with $\epsilon_{pp} = (0.0,\ 0.5,\ 1.0)\ k_B T$ and ideal polymers (with no excluded-volume and no cohesion).

[6] R. Avinery, M. Kornreich, and R. Beck, Universal and efficient entropy estimation using a compression algorithm, arXiv (2018), arXiv:1709.10164.

[7] J. Seward, Bzip2 and libbzip2: Version 1.0.5 A program and library for data compression http://www.bzip.org/ (2007).

[8] G. Strobl, *Phys. Polym. Concepts Underst. Their Struct. Behav.* (Springer, Berlin, 2007).

[9] L. E. Kapinos, R. L. Schoch, R. S. Wagner, K. D. Schleicher, and R. Y. H. Lim, Karyopherin-centric control of nuclear pores based on molecular occupancy and kinetic analysis of multivalent binding with FG nucleoporins, Biophys. J. **106**, 1751 (2014).

[10] R. Simpson, *J. Proteome Res. Proteins and Proteomics: A Laboratory Manual* (Cold Spring Harbour Laboratory Press, U.S., 2004).

[11] J. Yamada, J. L. Phillips, S. Patel, G. Goldfien, A. Calestagne-Morelli, H. Huang, R. Reza, J. Acheson, V. V. Krishnan, S. Newsam, A. Gopinathan, E. Y. Lau, M. E. Colvin, V. N. Uversky, and M. F. Rexach, A Bimodal Distribution of Two Distinct Categories of Intrinsically Disordered Structures with Separate Functions in FG Nucleoporins, Mol. Cell. Proteomics **9**, 2205 (2010).

[12] R. S. Wagner, L. E. Kapinos, N. J. Marshall, M. Stewart, and R. Y. H. Lim, Promiscuous binding of karyopherin$\beta$1 modulates FG nucleoporin barrier function and expedites NTF2 transport kinetics, Biophys. J. **108**, 918 (2015).

[13] D. P. Denning, V. Uversky, S. S. Patel, A. L. Fink, and M. Rexach, The Saccharomyces cerevisiae nucleoporin Nup2p is a natively unfolded protein, J. Biol. Chem. **277**, 33447 (2002).
